# Biofilm Patterning Reveals the Functional Contributions of Periplasmic Cytochromes to the Electrochemical Activity of *Shewanella oneidensis*

**DOI:** 10.64898/2026.01.07.698206

**Authors:** Fengjie Zhao, Christina M. Niman, Marko S. Chavez, Joshua T. Atkinson, Bridget E. Conley, Jeffrey A. Gralnick, James Q. Boedicker, Mohamed Y. El-Naggar

**Author notes:** Correspondence to: James Q. Boedicker, Mohamed Y. El-Naggar. Fengjie Zhao and Christina M. Niman contributed equally to this work.

## Abstract

*Shewanella oneidensis* MR-1 is a model electroactive bacterium whose extracellular electron transfer (EET) pathway includes a sequential network of *c*-type cytochromes that span the inner membrane, periplasm, and outer membrane. While electrochemical studies revealed the critical role of outer-membrane cytochromes in mediating both outward EET from cells to external surfaces and lateral biofilm conduction across cells, the specific functional role of the periplasmic cytochromes in these processes is less understood. Dissecting the contributions of the periplasmic components has been challenged by the complexity of the periplasmic cytochrome network and the inherent variability of native biofilms, which confounds the electrochemical comparison of cytochrome mutants. Here, we overcome these limitations with a synthetic biology approach combining targeted deletion of genes encoding key periplasmic cytochromes with light-induced biofilm patterning to create uniform, geometrically defined biofilms on electrodes for robust electrochemical comparisons. Voltammetric measurements of patterned *S. oneidensis* mutant biofilms confirmed the essential role of periplasmic cytochromes in facilitating outward EET, a contribution that becomes apparent when flavins are present to accelerate interfacial electron transfer between outer-membrane cytochromes and the electrode. In contrast to this crucial role in routing outward EET across the periplasm, electrochemical gating measurements of lateral biofilm conductivity revealed that the periplasmic cytochromes do not contribute to long-distance electron transport along cellular layers bridging electrodes. These findings provide new insights into the role of periplasmic cytochromes in *S. oneidensis*, and distinguish their contributions to routing outward EET across the cell envelope versus biofilm conductivity.

**Importance:** Microbes capable of extracellular electron transfer (EET) are central to global biogeochemical cycles and emerging bioelectrochemical technologies. In the important model EET bacterium *Shewanella oneidensis* MR-1, the outer-membrane components that interface with external surfaces are well-characterized. However, the functional role of the periplasmic components linking the inner and outer membranes has remained obscured by the complex network of multiple cytochromes and biofilm heterogeneity limiting precise comparisons across mutants. By combining light-induced biofilm patterning with electrochemical analysis, we successfully revealed the specific contributions of periplasmic cytochromes: these components are essential for facilitating the outward EET across the cell envelope but do not impact lateral long-distance electron transport across the biofilm. The results refine our understanding of extracellular respiration and provide design rules for engineering living electronic materials.

## Introduction

Electroactive bacteria can use extracellular electron transfer (EET) pathways to respire solid surfaces such as minerals and electrodes in anoxic environments for energy gain (1, 2). *Shewanella oneidensis* and *Geobacter metallireducens* were the two initially discovered electroactive bacteria, and the EET pathways of these two model electroactive organisms have been well studied (1–3). The reduction of extracellular electron acceptors by *S. oneidensis* requires an outward EET pathway from the inner membrane-associated cytochrome CymA (4), across the ∼23.5 nm wide periplasm (5), to the outer membrane-associated multiheme cytochrome-porin complex MtrCAB (6–8). Electrons are subsequently transferred to external electron acceptors either indirectly via soluble flavins acting as redox shuttles or directly through contact with cell surface cytochromes, flavin-cytochrome complexes, and micrometer-scale membrane extensions containing cytochromes that extend the reach of cells (9–12).

By analyzing interactions between cells and electrodes, biofilm electrochemistry provides crucial insight into the distinct modalities of electron flow in biofilms, namely outward EET and biofilm conduction. Outward EET describes the abovementioned net electron flux from cellular metabolism, across the cell envelope, to the electrode surface. Lateral biofilm conduction, on the other hand, refers to the long-distance electron transport process where electrons propagate across the biofilm (cell-to-cell) to bridge micrometer-scale gaps, even in the absence of metabolic turnover. In the case of outward EET, the contribution of outer-membrane components has been extensively studied with electrochemical measurements of mutants lacking outer-membrane cytochromes (10, 13). However, there has been no similar detailed electrochemical characterization of mutants deficient in periplasmic cytochromes to characterize their impact on outward EET to electrodes. In the context of biofilm conduction, recent electrochemical gating measurements of *S. oneidensis* biofilms bridging interdigitated electrodes showed the crucial role of outer-membrane cytochromes in mediating long-distance electron transport along and across cells (14, 15), but the extent to which periplasmic cytochromes participate in this micrometer-scale conduction network remains an open question (16).

Previous works based on protein structure modeling and cytochrome interactions have assumed that periplasmic cytochromes diffusively link the inner-membrane cytochrome CymA and outer-membrane cytochrome MtrA to allow electron transport across the periplasm in *S. oneidensis* (17–19). Studies have shown that periplasmic cytochromes CctA and FccA contribute to the reduction of soluble electron acceptors (such as Fe^3+^) (20–22). Additionally, overexpression of CctA has been shown to increase current production and power density during electrochemical measurements (21, 22). However, there has not yet been a systematic study of the contributions of *S. oneidensis* periplasmic cytochromes during electron transfer to external electrodes or during lateral biofilm conduction across electrodes. The diversity of the cytochromes present in the *S. oneidensis* periplasm has complicated efforts to fully understand their roles in EET. Recently, the complexity of the periplasmic cytochrome network in the model electroactive microorganism *Geobacter sulfurreducens* has been studied through gene deletion, metal reduction assays, and electrochemical measurements (23). The results demonstrated a lack of specificity among different periplasmic cytochromes during electron transfer, and none were required when cells utilized electrodes (23). These findings further raise the question of the role that periplasmic cytochromes play in *S. oneidensis* EET, the answer to which will deepen our understanding of EET mechanisms.

Electroactive bacteria are able to colonize electrode surfaces and form living conductive biofilms (24). Electrochemical techniques, such as cyclic voltammetry and electrochemical gating, are used to characterize conductive biofilms, as they can uncover underlying electron transport mechanisms and extract material properties (14, 15, 25). To study the role of periplasmic cytochromes in *S. oneidensis* EET, it is necessary to construct periplasmic cytochrome deficient *S. oneidensis* mutants and perform electrochemical measurements to compare the electrochemical activity of mutant and wild type strains. Over the past decade, synthetic biology and genome editing tools have been developed in *S. oneidensis* to delete the cytochrome genes or regulate gene expression to probe how certain cytochromes participate in electron transfer (26–29). However, unlike *G. sulfurreducens*, native *S. oneidensis* biofilms form patchy, non-uniform structures on electrodes (14); this introduces significant cross-electrode variability, confounding the subtle electrochemical phenotypic differences between cytochrome mutants. To address this issue, we recently developed a lithographic strategy to pattern *S. oneidensis* biofilms by using the light-induced genetic circuit pDawn to control the expression of cell aggregation proteins CdrAB (30). This system ensures the formation of thick, uniform biofilms with a defined geometry on electrodes (30), thereby facilitating consistent and reliable electrochemical comparisons of biofilms. In addition, this approach allowed quantification of the intrinsic lateral biofilm conductivity (30).

In this work, we combined synthetic biology and electrochemical techniques to study the role periplasmic cytochromes play in both *S. oneidensis* outward EET to electrodes and lateral biofilm conductivity. Two mutants, Δ*cctA*Δ*fccA* and Δ*cctA*Δ*fccA*Δ*nrfA*, lacking genes encoding the most prominent periplasmic cytochromes (20) were constructed and used in this study. We measured the soluble iron reduction capabilities of these mutants and the wild type. With this validation, we then used cyclic voltammetry and electrochemical gating to investigate the electrochemical activity and conductive properties of patterned biofilms of our mutants. Our results indicate that periplasmic cytochromes participate in the electron transfer to electrodes (outward flow of electrons from the cell), but do not contribute to the lateral biofilm conductivity. These findings deepen our understanding of the mechanisms *S. oneidensis* implements to transport electrons across its periplasm, offering insights into developing strategies to improve EET efficiency of living biofilms.

## Results

### Iron reduction assays validate that periplasmic cytochromes contribute to the reduction of soluble electron acceptors

A previous study showed that the periplasmic cytochromes CctA and FccA (Fig. 1) are involved in the reduction of soluble electron acceptors, such as ferric iron, DMSO, and nitrate (20). We first quantified soluble iron reduction capabilities of the wild type and our periplasmic cytochrome deficient mutants using a ferrozine assay. The results show that the double mutant Δ*cctA*Δ*fccA* exhibits a significant delay in iron reduction compared to the wild type (WT) (Fig. 2). Deleting the gene *nrfA* (Fig. 1), which encodes a periplasmic nitrite reductase (31), further delays soluble iron reduction in the triple mutant Δ*cctA*Δ*fccA*Δ*nrfA*. This suggests that NrfA also contributes to the reduction of soluble iron (Fig. 2). After the lag phase, the double and triple mutants begin to reduce soluble iron and eventually produce amounts of Fe^2+^ equivalent to those of the WT (Fig. 2). The times required for the double and triple mutants to fully reduce the soluble iron are around 3-fold longer than that of the WT. We checked the cytochrome expression of the triple mutant Δ*cctA*Δ*fccA*Δ*nrfA* at different time points during the iron reduction assay. MtrA abundance appeared to increase after the lag phase (Fig. S1), suggesting that MtrA may function both as part of the MtrCAB conduit and independently to facilitate periplasmic electron transfer. These results are consistent with a previous study (20), which reported a long lag phase in the growth of Δ*cctA*Δ*fccA* with iron citrate as the electron acceptor, along with higher MtrA expression after the lag phase. In summary, our results indicate that the periplasmic cytochromes CctA, FccA, and NrfA participate in *S. oneidensis* soluble iron reduction.

**Figure 1.**
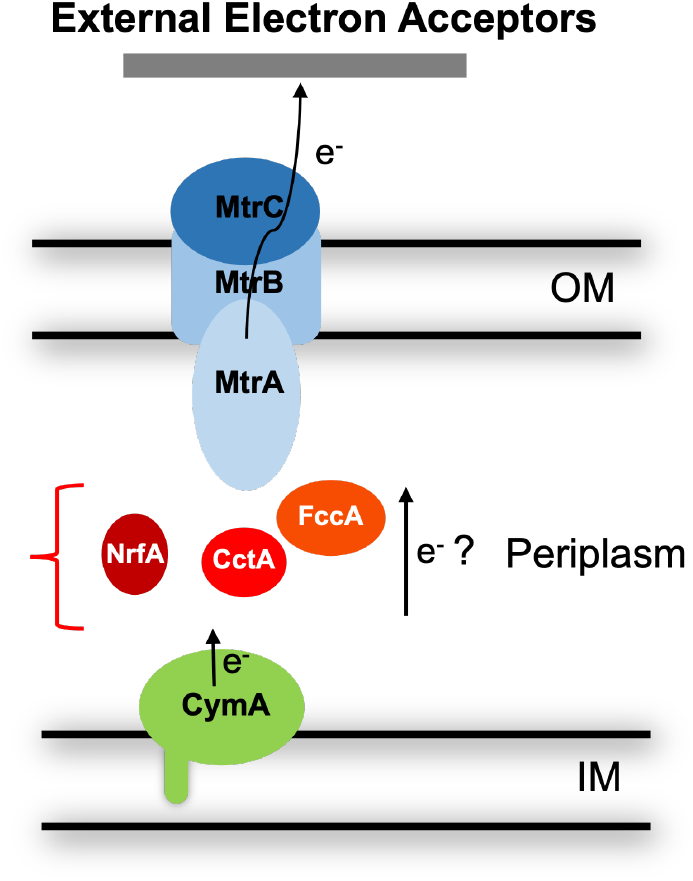
Schematic diagram of the *S. oneidensis c*-type cytochrome network. CctA, FccA, and NrfA are the most prominent cytochromes located in the periplasm of *S. oneidensis* (20). CctA: a small tetraheme cytochrome; FccA: a tetraheme cytochrome and fumarate reductase; NrfA: a pentaheme cytochrome and nitrite reductase; MtrCAB: outer-membrane cytochrome complex; CymA: inner-membrane cytochrome. IM: Inner membrane; OM: Outer membrane.

**Figure 2.**
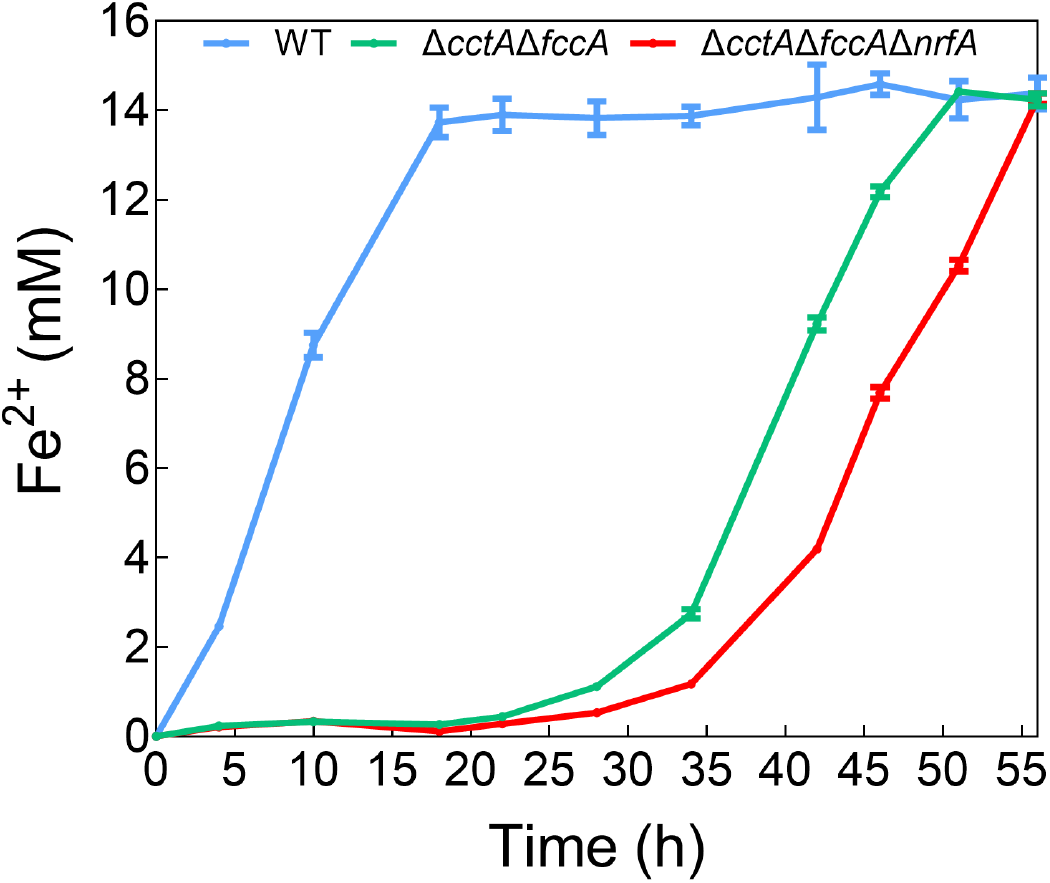
Iron reduction capabilities of *S. oneidensis* wild type and periplasmic cytochrome deficient mutants. WT: wild type strain. The data (mean ± SD) were obtained from triplicate measurements.

### Biofilm patterning enables uniform biofilm formation of periplasmic cytochrome deficient mutants

Next, the contribution of periplasmic cytochromes to electron transfer between *S. oneidensis* biofilms and electrodes was analyzed. Reliably comparing biofilm EET activity across *S. oneidensis* strains and mutants is often hindered by their patchy and nonuniform coverage on electrodes. To overcome this limitation, we utilized a genetic circuit to improve *S. oneidensis* biofilm formation (30). This circuit enables light-regulated production of the cell aggregation proteins CdrAB, which leads to the formation of thicker and more uniform biofilms with defined geometries. We transformed the plasmid pDawn-CdrAB, containing the biofilm patterning circuit, into the *S. oneidensis* strains used for electrochemical measurements. These new strains were named Δ*cctA*Δ*fccA*+CdrAB and Δ*cctA*Δ*fccA*Δ*nrfA*+CdrAB. The wild type transformed with the pDawn-CdrAB plasmid was named WT+CdrAB.

To verify the compatibility of the biofilm patterning construct in the context of our periplasmic deletion strains, a cell-cell aggregation assay was performed. Aggregation of cells following light-induced expression of CdrAB suggests that the strains possess improved biofilm formation capabilities (Fig. 3A). The aggregation index, which measures the reduction of the culture turbidity at the top of the culture tube after settling, shows higher values of WT+CdrAB, Δ*cctA*Δ*fccA*+CdrAB and Δ*cctA*Δ*fccA*Δ*nrfA*+CdrAB strains than that of the WT strain (Fig. 3B). These results indicate significant cell-cell adhesion in Δ*cctA*Δ*fccA*+CdrAB and Δ*cctA*Δ*fccA*Δ*nrfA*+CdrAB strains after illumination.

**Figure 3.**
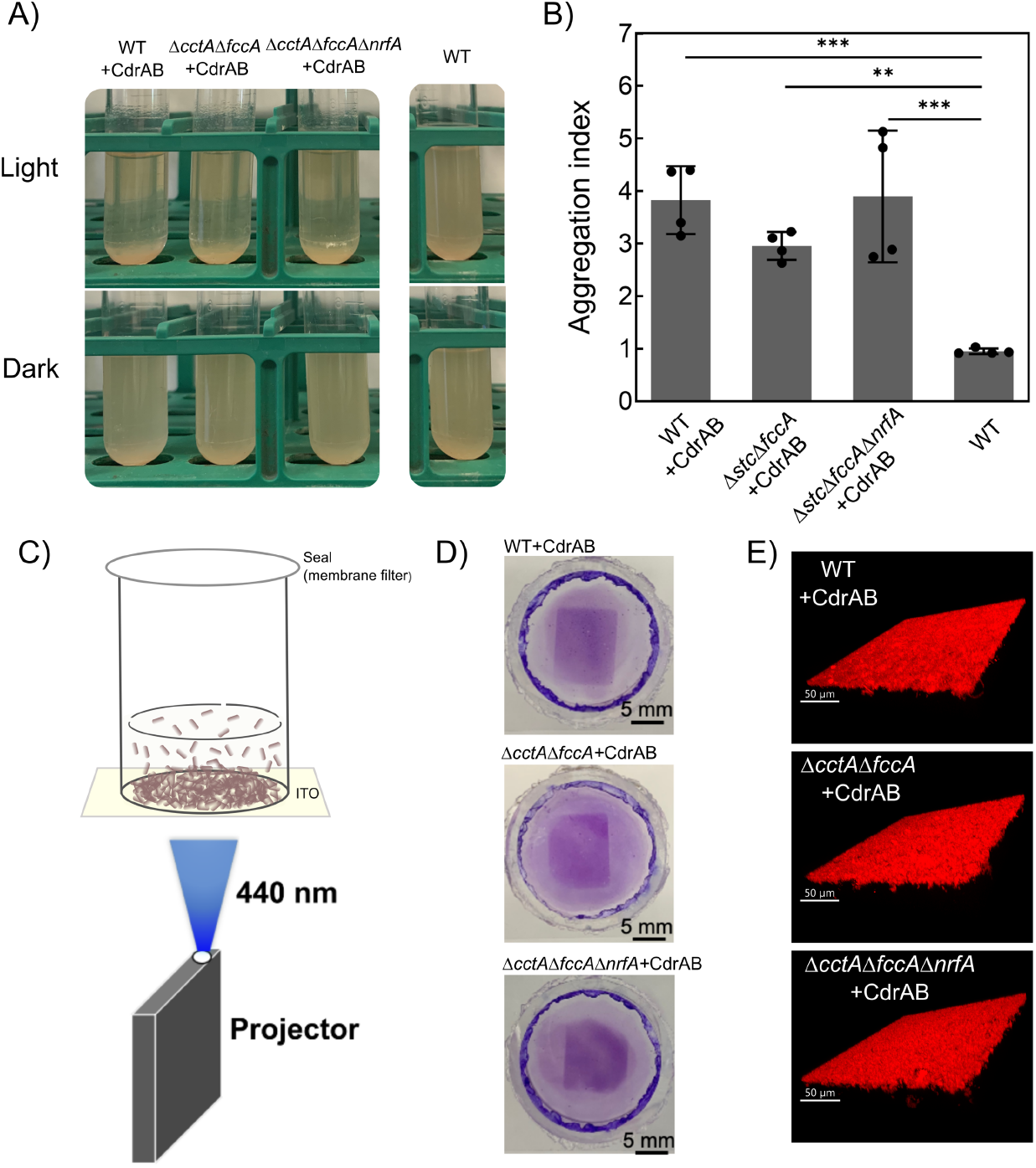
Cell-cell aggregation assay and biofilm patterning of different *S. oneidensis* strains. Cell-cell aggregation assay (A) and aggregation index measurements (B) of WT (wild type), WT+CdrAB, Δ*cctA*Δ*fccA*+CdrAB, and Δ*cctA*Δ*fccA*Δ*nrfA*+CdrAB. For the aggregation index, data (mean ± SD) were obtained from four biological replicates. *p* = 0.0003 for WT+CdrAB vs. WT, *p* = 0.0051 for Δ*cctA*Δ*fccA*+CdrAB vs. WT, and *p* = 0.0002 for Δ*cctA*Δ*fccA*Δ*nrfA*+CdrAB vs. WT (one-way ANOVA with Dunnett’s multiple comparisons test). Significance is indicated as \*\*\**p* < 0.001 and \*\**p* < 0.01. (C) Schematic diagram of light-induced biofilm patterning on an ITO electrode in our custom glass tube culturing vessel using a portable LED projector. (D) Crystal violet staining of patterned biofilms with defined geometry. (E) Confocal microscope images of patterned biofilms.

To facilitate electrochemical measurements, we also demonstrated biofilm patterning of these strains on the surface of transparent electrodes. Cells were cultured in custom vessels made by vertically attaching a glass tube to a conductive, planar indium tin oxide (ITO) coated glass coverslip (Fig. 3C). Rectangular patterns of blue light were projected onto the bottom of the ITO coverslip from below using a portable LED projector (Fig. 3C). The results show clear rectangular patterns of biofilms after staining with crystal violet (Fig. 3D). We also imaged the patterned biofilms via confocal microscopy which shows dense and uniform biofilms with a thickness of ∼ 10 μm (Fig. 3E and S2) (30). When our custom vessel was used ahead of electrochemical measurements, a thin copper wire was electrically connected to the ITO coverslip before cell culturing and patterning. After patterning onto the wired ITO coverslip, reference and counter electrodes were added to the vessel, allowing it to serve as a bioreactor for electrochemical measurements (Fig. 4A).

**Figure 4.**
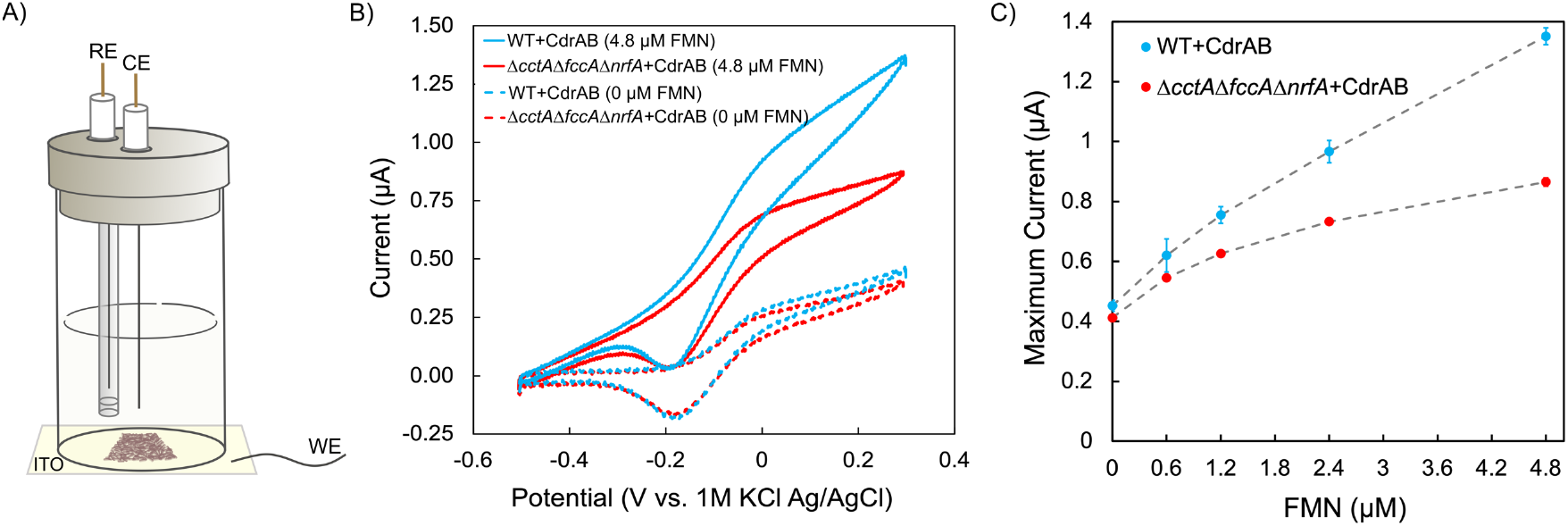
Electrochemical measurements of patterned WT+CdrAB and *ΔcctAΔfccAΔnrfA*+CdrAB biofilms on transparent ITO electrodes. (A) Schematic diagram of the bioreactor with patterned biofilms for electrochemical measurements. RE: reference electrode; CE: counter electrode; WE: working electrode. (B) Representative cyclic voltammograms of light patterned biofilms of WT+CdrAB and *ΔcctAΔfccAΔnrfA*+CdrAB with 0 µM and 4.8 µM flavin mononucleotide (FMN) added. (C) Effect of adding increasing concentrations of FMN on the maximum current of light patterned biofilms of WT+CdrAB and *ΔcctAΔfccAΔnrfA*+CdrAB. The data (mean ± SD) were obtained from two biological replicates.

### Periplasmic cytochromes participate in EET to electrodes

Biofilm patterning enables controlled biofilm formation, enabling more reproducible electrochemical measurements compared to native biofilms (Fig. S2) (30). This allows for the direct comparison of electrochemical results among different mutants and reactors. To study whether periplasmic cytochromes contribute to electron transfer to an electrode, we performed cyclic voltammetry measurements on patterned biofilms of WT+CdrAB, Δ*cctA*Δ*fccA*+CdrAB, and Δ*cctA*Δ*fccA*Δ*nrfA*+CdrAB on the ITO glass coverslip with a defined geometry of 1.2 cm x 1.0 cm (Fig. 4A). The voltammograms revealed that the current generation of Δ*cctA*Δ*fccA*+CdrAB and Δ*cctA*Δ*fccA*Δ*nrfA*+CdrAB was nearly identical to the WT+CdrAB strain (Fig. S3A), and that all three strains have similar maximum current densities (Fig. S3B).

The similar current production by all three strains was surprising because the periplasmic cytochrome deficient mutants show clear phenotypes for delayed iron reduction capability, but no obvious phenotype for electrode reduction. We reasoned that this could be due to electron transfer through the *S. oneidensis* periplasmic space not being a rate-limiting step under our experimental conditions. Previous experimental and computational studies have shown that the efficiency of interfacial (*i*.*e*., cell-to-electrode) electron transport was enhanced when flavins interact with flavin-binding sites on the outer-membrane cytochromes or act as extracellular redox shuttles (9, 32–34), suggesting that the interfacial step may be rate-limiting in the absence of flavins. In our electrochemical measurements, repeated washing and replacement with fresh, flavin-free minimal medium removed extracellular flavins, such that the final interfacial electron transfer likely became rate-limiting. To examine how enhancing this interfacial step may affect our investigation, we conducted additional electrochemical measurements for the patterned biofilms of WT+CdrAB and Δ*cctA*Δ*fccA*Δ*nrfA*+CdrAB, in which different concentrations of flavins were exogenously added. The representative cyclic voltammograms show that the currents produced by Δ*cctA*Δ*fccA*Δ*nrfA*+CdrAB are lower than those of the WT+CdrAB when 4.8 µM flavin mononucleotide (FMN) is added (Fig. 4B), while the voltammograms of WT+CdrAB and Δ*cctA*Δ*fccA*Δ*nrfA*+CdrAB overlap without FMN addition (Fig. 4B). The differences in the voltammograms between WT+CdrAB and Δ*cctA*Δ*fccA*Δ*nrfA*+CdrAB become more pronounced as more FMN is added (Fig. S4). Figure 4C shows the maximum currents from the WT+CdrAB and Δ*cctA*Δ*fccA*Δ*nrfA*+CdrAB voltammograms as a function of FMN addition (Fig. 4C). The results indicate that there is no obvious difference between the maximum currents of WT+CdrAB and Δ*cctA*Δ*fccA*Δ*nrfA*+CdrAB without exogenous FMN addition (Fig. 4C). However, a lower maximum current is observed from Δ*cctA*Δ*fccA*Δ*nrfA*+CdrAB, as compared to WT+CdrAB, with the first addition of 0.6 µM FMN. Increasing differences between the maximum currents of WT+CdrAB and Δ*cctA*Δ*fccA*Δ*nrfA*+CdrAB arise with each successive addition of FMN (Fig. 4C).

Taken collectively, we combined light-induced biofilm patterning and electrochemical measurements to study the electrode reduction phenotype of periplasmic cytochrome deficient mutants. These results indicate that the periplasmic cytochromes participate in shuttling electrons through the periplasmic space during electrode reduction. This contribution is most apparent when flavins are added to eliminate interfacial cell-electrode electron transport limitations.

### Periplasmic cytochromes do not impact lateral biofilm conductivity

In addition to interfacial electron transfer to electrodes, electroactive biofilms can act as living electronic materials, allowing for long-distance electron transport across neighboring cells on the micrometer-scale (35). Previous studies demonstrated a crucial role for cell surface multiheme cytochromes in this biofilm conduction, but the extent to which periplasmic cytochromes may participate in the conduction mechanism is yet to be investigated (16). To determine the role of periplasmic cytochromes in biofilm conductivity, we patterned WT+CdrAB and Δ*cctA*Δ*fccA*Δ*nrfA*+CdrAB biofilms with a defined geometry (1.2 cm x 1.0 cm) onto custom ITO interdigitated array (IDA) electrodes by culturing the cells in custom vessels (Fig. 3C) with a transparent ITO IDA electrode serving as the bottom. The ITO IDA electrode contains two electrically isolated conductive regions that function as source and drain electrodes with an interspersed nonconductive region (Fig. 5A). To measure the conduction current through the patterned biofilms bridging the source and drain electrodes, electrochemical gating measurements were performed after biofilm patterning. When performing electrochemical gating measurements, the potentials at each working electrode are swept across a defined range while a potential offset (*V*_*SD*_) is maintained between the source and drain working electrodes to serve as a driving force for conduction. Figure 5B shows representative conduction current data for WT+CdrAB and Δ*cctA*Δ*fccA*Δ*nrfA*+CdrAB biofilms patterned onto IDAs with 20 µm insulating gaps. By plotting the conduction current (*I*_*cond*_) *vs* the gate potential (*E*_*G*_, average potential of the working electrodes during the gating scan), we observe a conduction peak centered at the formal potential of *S. oneidensis* cytochromes for both WT+CdrAB and Δ*cctA*Δ*fccA*Δ*nrfA*+CdrAB (Fig. 5B), consistent with a redox conduction mechanism (15). Both WT+CdrAB and Δ*cctA*Δ*fccA*Δ*nrfA*+CdrAB show overlapping conduction currents and statistically indistinguishable peak magnitudes (Fig. 5B and S5). With the known peak conduction current, defined biofilm geometry, and uniform biofilm thickness (∼10 µm for patterned biofilms), we calculated the conductivity of the WT+CdrAB and Δ*cctA*Δ*fccA*Δ*nrfA*+CdrAB biofilms to be 6.4 ± 0.9 nS/cm and 6.0 ± 0.9 nS/cm, respectively (Fig. 5C). No significant difference in conductivity was observed between the two strains. These results indicate that the periplasmic cytochromes do not contribute to lateral biofilm conductivity.

**Figure 5.**
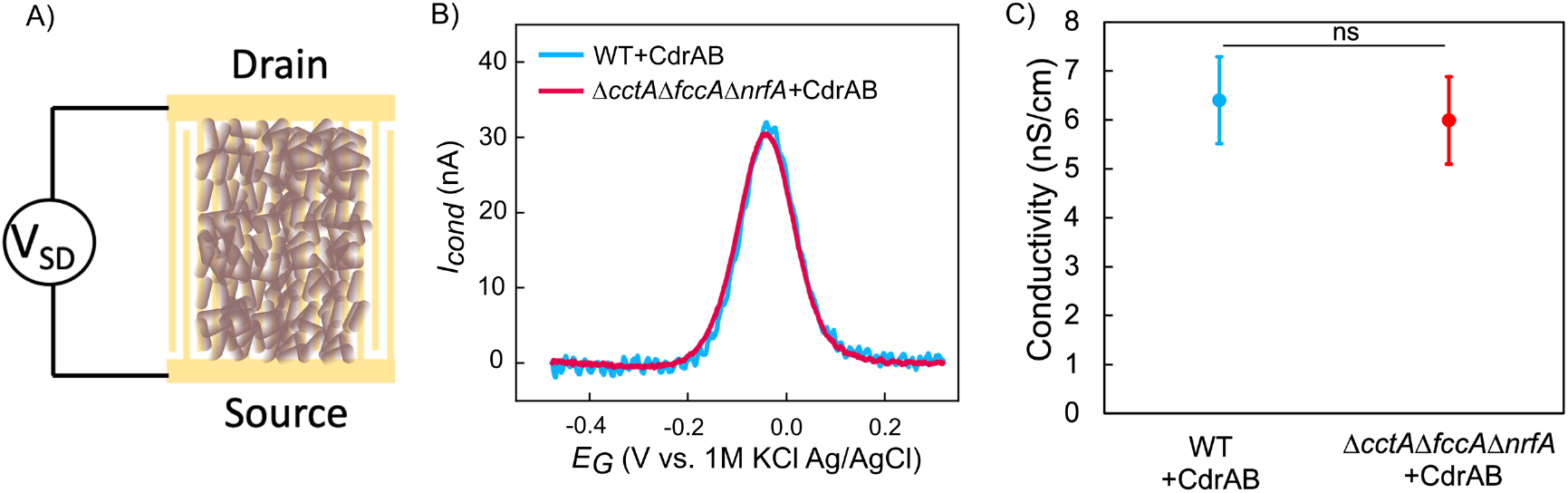
Conductivity measurements of WT+ CdrAB and *ΔcctAΔfccAΔnrfA*+CdrAB patterned biofilms on ITO interdigitated array (IDA) electrodes by electrochemical gating. (A) Schematic diagram of the electrode configuration used in electrochemical gating measurement of patterned biofilms on IDA electrodes. (B) Representative conduction currents of patterned WT+CdrAB and Δ*cctA*Δ*fccA*Δ*nrfA*+CdrAB biofilms on IDAs. (C) Conductivities of patterned WT+CdrAB and Δ*cctA*Δ*fccA*Δ*nrfA*+CdrAB biofilms extracted from the peak conduction currents and known biofilm and electrode geometries. No significant difference in conductivity was observed between patterned WT+CdrAB biofilms and Δ*cctA*Δ*fccA*Δ*nrfA*+CdrAB biofilms (two-tailed unpaired *t*-test, *p* = 0.6021). Data are presented as mean ± SD from three biological replicates. Significance is indicated as ns (not significant, *p* > 0.05).

To further confirm that our conductivity measurements are performed under conditions where homogeneous electron transport (conduction within the biofilm) is rate-limiting, rather than heterogeneous EET (biofilm-electrode injection/ejection), we performed scan rate dependent non-turnover cyclic voltammetry of patterned WT+CdrAB biofilms on IDAs (15). Figure S6 shows the representative voltammograms collected at 8 different scan rates spanning from 1 to 200 mV/s with *V*_*SD*_ = 0.0 V. Under these conditions, the resulting current is solely due to the charging and discharging of redox cofactors in the biofilm. The scaling relationship between the current and the scan rate provides information as to whether the rate-limiting step for biofilm conduction is homogeneous (*i*.*e*., between cells) or heterogeneous (*i*.*e*., between cells and the electrode) electron transfer (15). The peak currents observed in the data increase with increasing scan rate and scale linearly with the square root of the scan rate (Fig. S6 Inset). The positions of the peaks do not change with increasing scan rate. These results show that the rate-limiting step of electron transport is diffusion within the biofilms, and not electron transfer at the electrode-biofilm interface. Therefore, the similar conductivity values of WT+CdrAB and Δ*cctA*Δ*fccA*Δ*nrfA*+CdrAB biofilms measured from our electrochemical gating experiments reflect nearly identical electron transport within the biofilm.

## Discussion

Periplasmic cytochromes are expected to mediate electron flux between the inner and outer membranes in *S. oneidensis* (17–19), thereby facilitating outward EET to external electron acceptors. However, their role has so far been directly assessed only in reducing soluble electron acceptors. In this work, we studied the role of periplasmic cytochromes in soluble iron reduction, outward EET to electrodes, and lateral biofilm conduction across electrodes by combining synthetic biology strategies with electrochemical techniques. Outer-membrane cytochromes, such as MtrC, were previously shown to be important for soluble iron reduction (36). Since periplasmic cytochromes CctA and FccA are thought to carry electrons to the outer-membrane MtrCAB complex, we first measured the iron citrate reduction capabilities of periplasmic cytochrome deficient mutants. Our results indicate that all periplasmic cytochromes (CctA, FccA and NrfA) play a role in iron reduction, but also that *S. oneidensis* possesses a mechanism to compensate for the activity of the missing cytochromes over time. These results are consistent with a previous study, which showed that while the double mutant Δ*cctA*Δ*fccA* had a lag phase in growth with soluble iron as the electron acceptor, it could catch up to the growth of wild type by upregulating the expression levels of MtrA and NrfA (20). In our work, MtrA also appeared to have increased expression in the triple mutant Δ*cctA*Δ*fccA*Δ*nrfA* after the lag phase during soluble iron reduction (Fig. S1). These results indicate a lack of periplasmic cytochrome specificity in soluble iron reduction and that *S. oneidensis* has alternative mechanisms to recover iron reduction in the absence of CctA, FccA and NrfA. It is interesting to note that a lack of specificity for periplasmic electron transfer has also been observed recently in another model EET organism, *Geobacter sulfurreducens*, where any of 6 triheme cytochromes supported comparable growth on soluble or insoluble metal (23). It appears that this general promiscuity in the periplasmic electron pathways may be a general feature of outward EET.

Our initial electrochemical measurements of patterned biofilms show that deletion of the periplasmic cytochromes has no significant effect on electrode reduction without the addition of flavins (Fig. S3). However, we propose that during electrochemical measurements, the periplasmic electron transport rate was masked by the slower interfacial transfer between outer-membrane cytochromes and the electrode. Previous studies have shown that the efficiency of this interfacial step is enhanced when flavins are present to interact with flavin-binding sites on the outer-membrane cytochromes or act as soluble redox shuttles (9, 32–34). Accordingly, under our experimental conditions in which extracellular flavins were washed out prior to biofilm electrochemistry, the final electron transfer between outer-membrane cytochromes and the electrode is rate-limiting, regardless of which periplasmic cytochromes are present. With flavin addition, the electron transfer through the periplasm can become rate-limiting, making the role of periplasmic cytochromes observable. Consistent with this, electrochemical measurements of patterned biofilms with the addition of flavins show significantly lower current production in Δ*cctA*Δ*fccA*Δ*nrfA*+CdrAB compared to WT+CdrAB (Fig. 4). These results indicate that periplasmic cytochromes play a role in routing electron transfer across the periplasm to outer-membrane components, and their contribution becomes evident under conditions where the terminal cell-electrode step is no longer rate-limiting. We thus conclude that, while periplasmic cytochromes play a role in electron transport across the periplasm during EET, this role does not appear to be specific to any individual cytochrome. *S. oneidensis* possesses sufficient electron transport machinery for moving electrons across the periplasm to meet the metabolic needs of the biofilm. Most likely, the deficiencies introduced by deleting genes encoding various periplasmic cytochromes are compensated by alternative periplasmic cytochromes (including MtrA), as recently observed in *Geobacter* (23).

Finally, the electrochemical gating results shed light on the role of the periplasmic cytochromes in biofilm conduction along/across cellular layers bridging electrodes. Our previous studies show good agreement between the measured *S. oneidensis* biofilm conductivity (∼4.5 nS/cm) from the patterned biofilms (30) and the calculated *S. oneidensis* biofilm conductivity (∼7 nS/cm) based on a collision-exchange model taking into account the dynamics of outer-membrane cytochromes (16). However, any possible contribution to this process from periplasmic components remained an open question. In this work, our measured conductivities show no significant difference between the WT+CdrAB biofilms (6.4 ± 0.9 nS/cm) and the Δ*cctA*Δ*fccA*Δ*nrfA*+CdrAB biofilms (6.0 ± 0.9 nS/cm). We further ruled out that slow heterogeneous EET (between the biofilm and interdigitated electrodes) may be rate-limiting in biofilm conduction, thereby obfuscating rate differences in homogeneous conduction (through the biofilm) when comparing WT+CdrAB and Δ*cctA*Δ*fccA*Δ*nrfA*+CdrAB. Our scan-rate dependent non-turnover cyclic voltammetry measurements confirm that the rate-limiting step of biofilm conduction is electron diffusion within the biofilm, and not injection/ejection at the electrode-biofilm interface. These results indicate that the periplasmic cytochromes CctA, FccA, and NrfA do not play a significant role in lateral biofilm conduction of the patterned *S. oneidensis* biofilms. Consistent with this conclusion, recent work by Wen et al. showed that deletion of periplasmic cytochromes had a negligible impact on the thermal activation energy (*E*_a_) of electron transport in *Shewanella* biofilms (37). Our findings support a model where lateral conductivity is mediated primarily by the outer-membrane cytochrome network, consistent with the previously proposed collision-exchange mechanism (16).

The results presented in this work provide new insights on how periplasmic cytochromes participate in electron transport both across the periplasm and within biofilms, which deepens our understanding of the EET mechanism in *S. oneidensis*. Our work also shows that controlling the biofilm formation of *S. oneidensis* on electrodes allows us to perform reliable comparisons of electrochemical measurements among different mutants and reactors, which is a promising strategy to quantitatively characterize EET mechanisms in other electroactive bacteria, especially those that form inconsistent biofilms.

## Materials and methods

### Bacterial strains, plasmids and media

The strains and plasmids used in this study are listed in Table S1. The *S. oneidensis* Δ*cctA*Δ*fccA*Δ*nrfA* (JG4347) strain was generated by deleting *nrfA* in strain Δ*cctA*Δ*fccA* (JG3107) (38) utilizing materials and methods described previously (39). The blue light-induced biofilm patterning plasmid pDawn-CdrAB was transformed into the Δ*cctA*Δ*fccA* and Δ*cctA*Δ*fccA*Δ*nrfA* to get the Δ*cctA*Δ*fccA*+CdrAB and Δ*cctA*Δ*fccA*Δ*nrfA*+CdrAB strains.

*S. oneidensis* strains were cultivated in lysogeny broth (LB) medium (10 g/L tryptone, 5 g/L yeast extract, and 5 g/L sodium chloride) or minimal medium (15.1 g/L PIPES buffer, 3.4 g/L sodium hydroxide, 1.5 g/L ammonium chloride, 0.1 g/L potassium chloride, 0.6 g/L sodium phosphate monobasic monohydrate, 3.36 g/L 60% (w/w) sodium lactate, 1 mL/L mineral solution, 1 mL/L vitamin solution, 10 mL/L amino acid solution, pH 7.0) (38). For plasmid maintenance, media were supplemented with spectinomycin (Spec, 100 μg/mL). For the iron reduction assays, the minimal medium was used with 20 mM iron (III) citrate as the electron acceptor. For cyclic voltammetry measurements, the minimal medium was used without the vitamin solution to exclude the contribution from flavins. For electrochemical gating measurements, the minimal medium was used without sodium lactate and the vitamin solution. The media used for the iron reduction assays and the electrochemical measurements were purged with nitrogen.

### Iron reduction assay

The ferrozine assay was used to measure Fe^2+^ - a byproduct of Fe^3+^ reduction of *S. oneidensis* strains. Cells were aerobically cultured in LB medium overnight. Cells were then collected by centrifuging (5840R, Eppendorf) at 3,234 x g, 4 °C for 15 min and washed with fresh minimal medium 2 times. Cells were inoculated into sealed serum bottles containing 100 mL of anaerobic minimal medium to an OD600 of 0.1. 20 mM ferric citrate was added into the anaerobic minimal medium as the electron acceptor. The samples were incubated at 30 °C and 200 rpm with an orbital throw of 2.5 cm. At different time points, samples were taken out from the serum bottles and 10 μL of each sample was added immediately to 90 μL of 1 M HCl in a 96-well plate followed by 100 μL of 0.1% ferrozine. After the samples were mixed well and allowed to sit for 10 min, the absorbance of the samples at 562 nm was determined with a plate reader (Infinite 200 PRO, Tecan). A standard curve of freshly made ferrous sulfate was used to determine the Fe^2+^ concentrations.

### Cell-cell aggregation assay and biofilm patterning

The cell-cell aggregation assay was performed as described previously (30). Briefly, late log phase LB cultures of *S. oneidensis* strains were transferred into (1%, v/v) minimal medium and incubated aerobically overnight at 30 °C and 200 rpm with an orbital throw of 2.5 cm under either blue light or dark conditions. Then, the cultures were left to rest for 30 min. The upper region of cultures in tubes were collected to measure the OD_top_. Then, the cultures were rigorously vortexed for 10 s and the OD_total_ was determined. 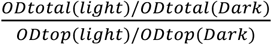 was used to calculate the aggregation index.

Biofilm patterning was performed as described in our previous work (30). The late log phase LB cultures (OD600 about 1−1.5) were diluted into fresh minimal medium to an OD600 of 0.01. 1 mL of the dilution was added to the bioreactor, which was made by attaching a 20 mm diameter and 2.5 cm tall glass tube onto a transparent working electrode (WE, ITO coated glass coverslip or custom ITO IDA). The glass tube was sealed with a microporous membrane filter (AirOtop Seals) and incubated at 30 °C aerobically. A portable smart projector (A5 Pro, Wowoto) was secured below the bioreactor in the incubator and 440 nm blue light patterns created in Microsoft PowerPoint were projected onto the underside of the bioreactor (Fig. 3C). After 16 h of patterning, medium in the bioreactor was discarded, and the patterned biofilms were washed 3 times, for 2 min each time, with minimal medium on an orbital shaker at 60 rpm to reduce the nonspecific patterned cells. The bioreactor with fresh minimal medium was then moved to the anaerobic chamber for electrochemical measurements of the patterned biofilms. To assess biofilm formation and patterning on the electrodes, crystal violet staining was performed either immediately after patterning (if no electrochemical measurements were conducted) or after the measurements (if they were needed).

### Cyclic voltammetry measurements

The cyclic voltammetry (CV) measurements were performed in an anaerobic chamber (Bactron 300, Sheldon Manufacturing, Inc.) with a 95:5 (N2/H2) atmosphere. After washing the patterned biofilms, the minimal medium in the bioreactor was exchanged for anoxic minimal medium without vitamin solution inside the anaerobic chamber before CV measurements. The CV measurements were conducted with a three-electrode reactor setup. PEEK plastic lids were used with the bioreactors along with platinum wire counter electrodes (CEs) and Ag/AgCl (1M KCl) reference electrodes (REs, CHI 111) (Fig. 4A). The PEEK plastic lids along with CEs and REs were not present during biofilm patterning and only added after biofilm patterning and washing once the bioreactors were in the anaerobic chamber. The CV measurements of patterned biofilms were performed from −0.5 to 0.3 V at 1 mV/s using a four-channel potentiostat (Admiral Instruments Squidstat). Three cycles were performed for each CV, and data from only the third cycle was presented in this work. All potentials reported in this work are vs Ag/AgCl (1 M KCl).

### Electrochemical gating measurements for analyzing biofilm conduction and conductivity

The custom indium tin oxide (ITO) interdigitated array (IDA) electrodes were designed and fabricated as described in our previous work (30). The interdigitated area (1.2 x 1 cm^2^) of the ITO IDA consisted of electrode fingers (12 mm long and 10 μm wide of each finger) and gaps (20 um between every two fingers). The electrochemical gating measurements were performed under non-turnover conditions (minimal media without sodium lactate and vitamin solution) and conducted under a 95:5 (N2/H2) atmosphere in an anaerobic chamber. Like the reactor setup used for our CV measurements, platinum wire counter electrodes (CEs), and Ag/AgCl (1 M KCl) reference electrodes (REs) were inserted into the reactors through the PEEK plastic lids. Electrochemical gating measurements were performed with a potentiostat (BioLogic VSP-300). The IDA source (*S*) and drain (*D*) working electrodes (WEs) were scanned at a rate of 1 mV/s from −0.5 to 0.3 V but with a fixed gating offset *V*_*SD*_ = 20 mV, where *V*_*SD*_ = *E*_*D*_ − *E*_*S*_ and where *E*_*D*_ and *E*_*S*_ are the potentials at each of the IDA WEs. Three electrochemical gating cycles were performed and data from only the third cycle was analyzed.

The conduction current *I*_*cond*_ can be extracted from the electrochemical gating measurements with the equation *I*_*cond*_ = (*I*_*D*_ − *I*_*S*_)/2, where *I*_*D*_ is the drain current and *I*_*S*_ is the source current (14).

The conductivity of the patterned biofilms was calculated based on the following equations (15) as described in our previous work (30).

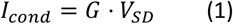

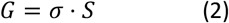

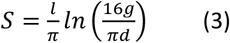

Equation 1 is used to determine biofilm conductance (*G*). Here *I*_*cond*_ is given by the peak conduction current and *V*_*SD*_ = 20 mV. In Equation 2, *G* is equal to the intrinsic conductivity of the biofilm (*σ*) multiplied by a geometric factor (*S*). For the case of uniform biofilms on an IDA with a thickness on the order of the size of the finger gaps, the geometric factor *S* is shown as Equation 3, based on a conformal mapping method. In equation 3, *l* is the length of the gap that snakes between the interdigitated fingers, *g* is the biofilm thickness, and *d* is the width of a finger gap.

### In situ confocal microscopy observations of patterned biofilms on electrodes and biofilm thickness analysis

After electrochemical measurements, the reactor media was discarded, and 2 mL of minimal medium containing FM 4-64FX membrane dye was added to the bioreactors to stain the biofilms for 30 mins. Then, the media was exchanged for 3% glutaraldehyde solution to fix the biofilms at 4 °C overnight. Before imaging the biofilms, the samples were washed with PBS three times. A Zeiss LSM 880 inverted microscope equipped with an argon laser and a 40× water objective lens was used to image the biofilms.

Imaris software was used to process confocal microscopy data and generate the 3D and cross-sectional biofilm images. Biofilm thickness measurements were performed utilizing Matlab by analyzing the cross-sectional confocal images as described in our previous work (30).

## Acknowledgements

This study was supported by the US Office of Naval Research Multidisciplinary University Research Initiative Grant No. N00014-18-1-2632. The fabrication of the IDA electrodes used in this work was performed at the San Diego Nanotechnology Infrastructure (SDNI) of UCSD, a member of the National Nanotechnology Coordinated Infrastructure, which is supported by the National Science Foundation (Grant ECCS-2025752). J.T.A. was supported by the NSF Postdoctoral Research Fellowships in Biology Program under grant no. 2010604.

