## Supplementary Information for "Biofilm Patterning Reveals the Functional Contributions of Periplasmic Cytochromes to the Electrochemical Activity of *Shewanella oneidensis*"

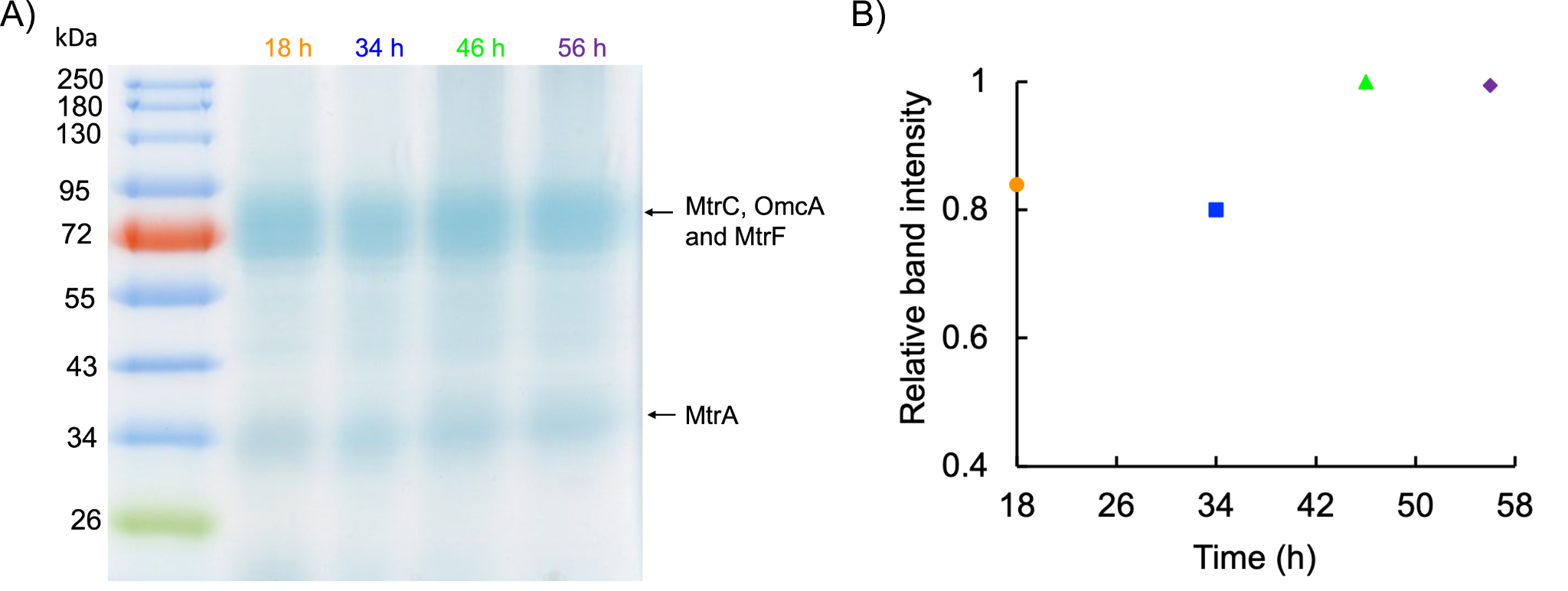

Figure S1. Cytochrome expression of the triple deficient mutant ∆*cctA*∆*fccA*∆*nrfA* at different time points of the iron reduction measurements. (A) TMBZ heme stain SDS-PAGE protein gel of ∆*cctA*∆*fccA*∆*nrfA* at different time points. (B) Band intensity of cytochrome MtrA at different time points. 18 h and 34 h are the time points during the lag phase of the iron reduction. 46 h and 56 h are the time points after the lag phase of the iron reduction.

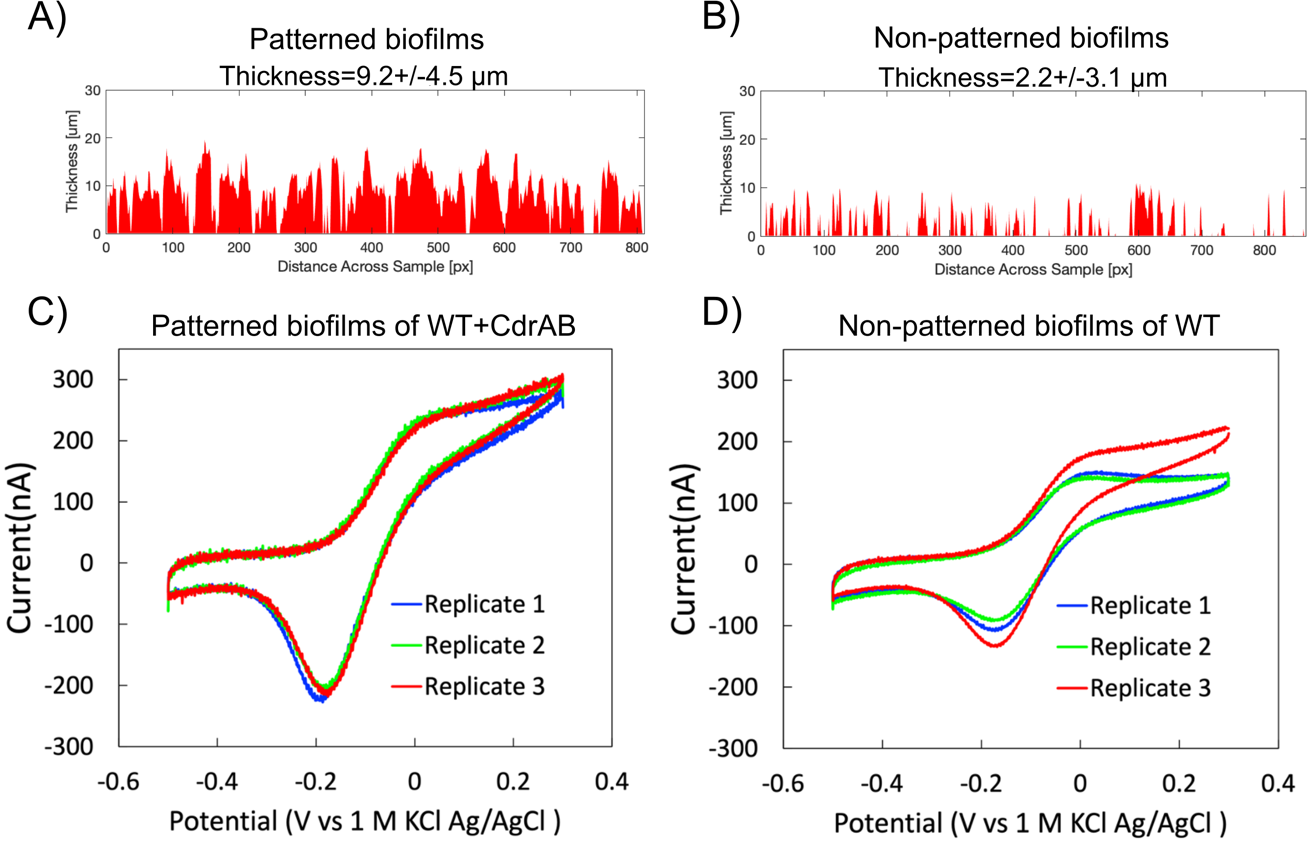

Figure S2. Biofilm thickness and cyclic voltammetry measurements of blue-light patterned biofilms and non-patterned native biofilms. (A) Biofilm thickness of patterned biofilms and non-patterned native biofilms measured from the cross-sectional confocal images based on the method used in our previous work (1). The patterned biofilms, with an average thickness of ∼ 10 μm, are more uniform and thicker than the non-patterned native biofilms with an average thickness of ∼ 2 μm. (B) Cyclic voltammetry measurements of patterned biofilms and non-patterned native biofilms. Patterned biofilms gave more consistent currents across different replicates than those of the non-patterned native biofilms.

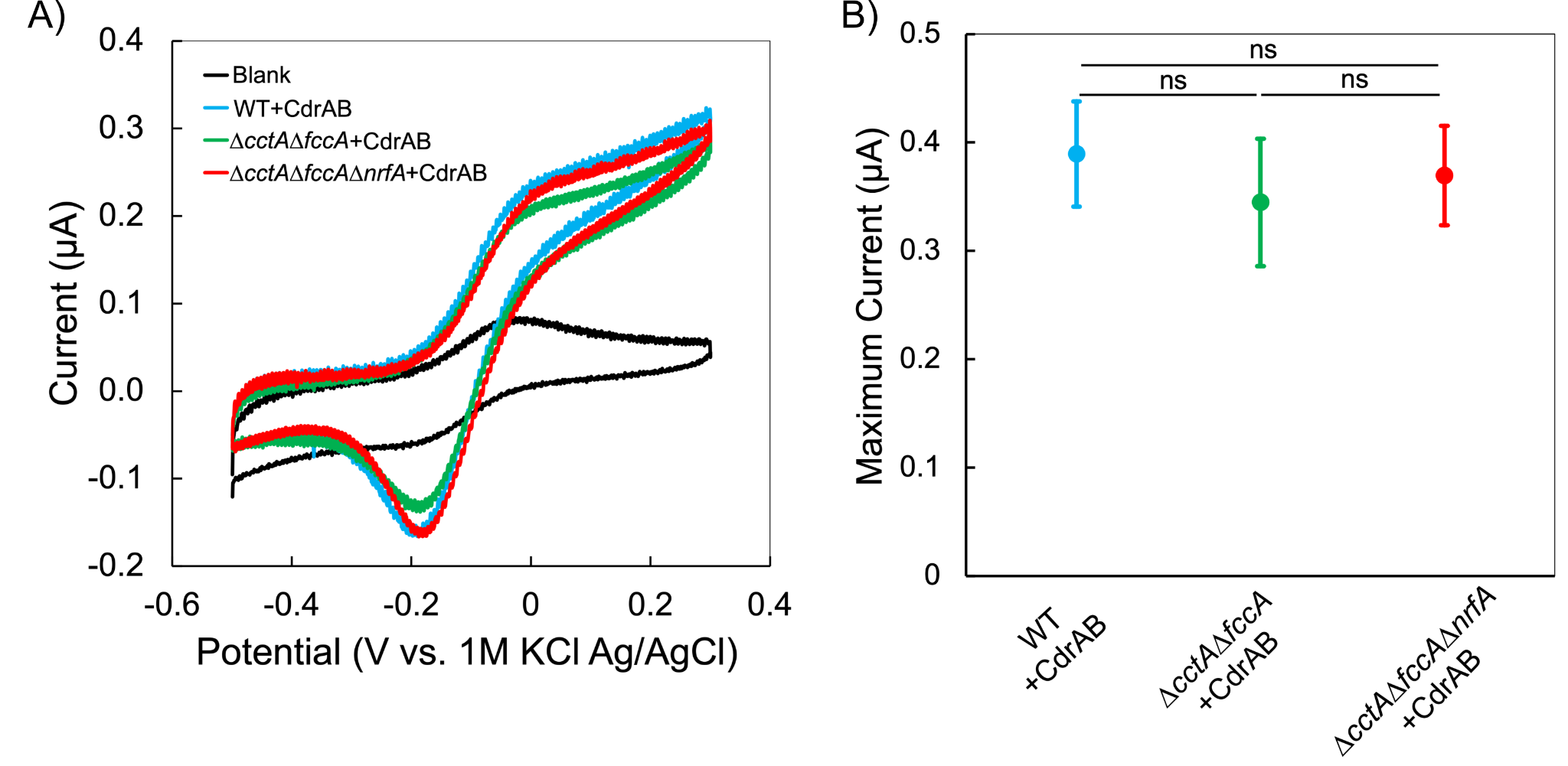

Figure S3. Cyclic voltammetry measurements of WT+CdrAB, ∆*cctA*∆*fccA*+CdrAB, and ∆*cctA*∆*fccA*∆*nrfA*+CdrAB patterned biofilms without adding flavins. (A) Representative cyclic voltammograms of blue light patterned biofilms of WT+CdrAB, ∆*cctA*∆*fccA*+CdrAB, and ∆*cctA*∆*fccA*∆*nrfA*+CdrAB. (B) The maximum currents from the cyclic voltammograms for each mutant. For quantitative comparison, the maximum current of each voltammogram was collected. The means and standard deviations were calculated from multiple biological replicates: 11 for the WT+CdrAB, 4 for the ∆*cctA*∆*fccA*+CdrAB, and 5 for the ∆*cctA*∆*fccA*∆*nrfA*+CdrAB. The maximum current is 0.39 ± 0.05 µA for the WT+CdrAB, 0.34 ± 0.06 µA for the ∆*cctA*∆*fccA*+CdrAB and 0.37 ± 0.05 µA for the ∆*cctA*∆*fccA*∆*nrfA*+CdrAB. No significant differences in maximum currents from the cyclic voltammograms were observed among the three patterned biofilms (two-tailed unpaired *t*-test, *p* = 0.1594 for WT+CdrAB vs. ∆*cctA*∆*fccA*+CdrAB, *p* = 0.4547 for WT+CdrAB vs. ∆*cctA*∆*fccA*∆*nrfA*+CdrAB, and *p* = 0.5003 for ∆*cctA*∆*fccA*+CdrAB vs. ∆*cctA*∆*fccA*∆*nrfA*+CdrAB). Significance is indicated as ns (not significant, *p* > 0.05).

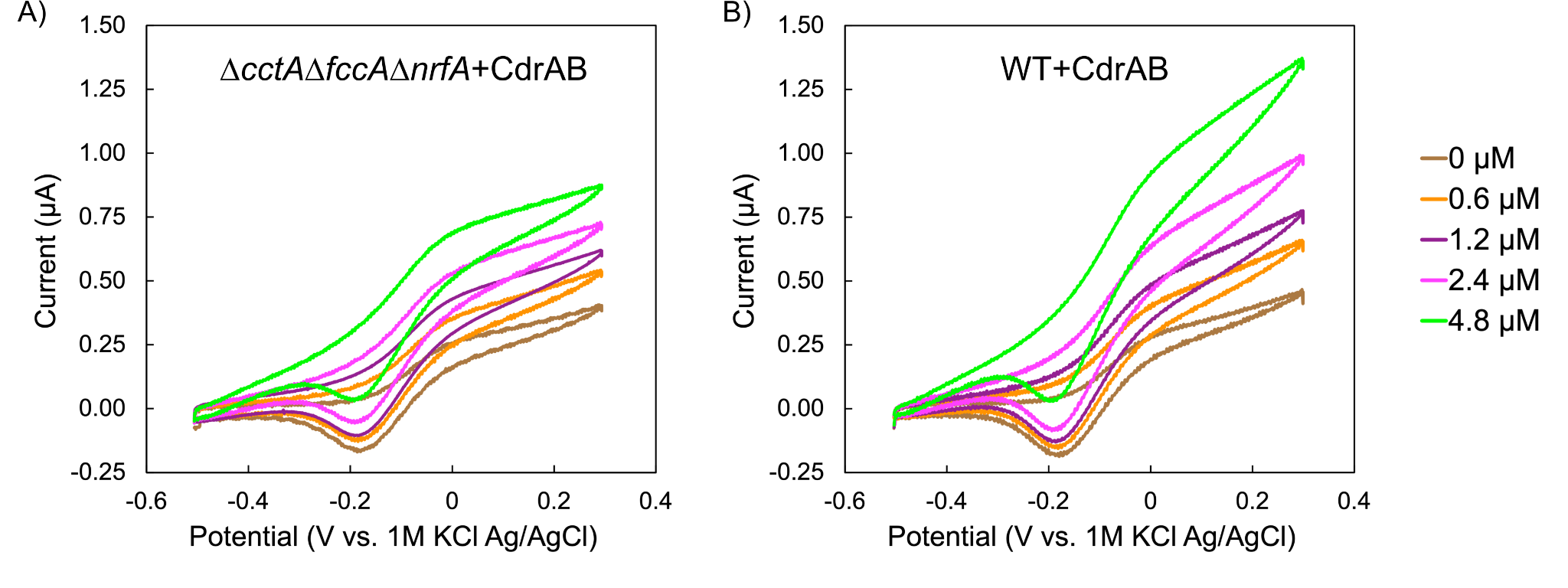

Figure S4. Representative cyclic voltammetry measurements of ∆*cctA*∆*fccA*∆*nrfA*+CdrAB (A) and WT+CdrAB (B) patterned biofilms with different concentrations of flavin mononucleotide (FMN) added.

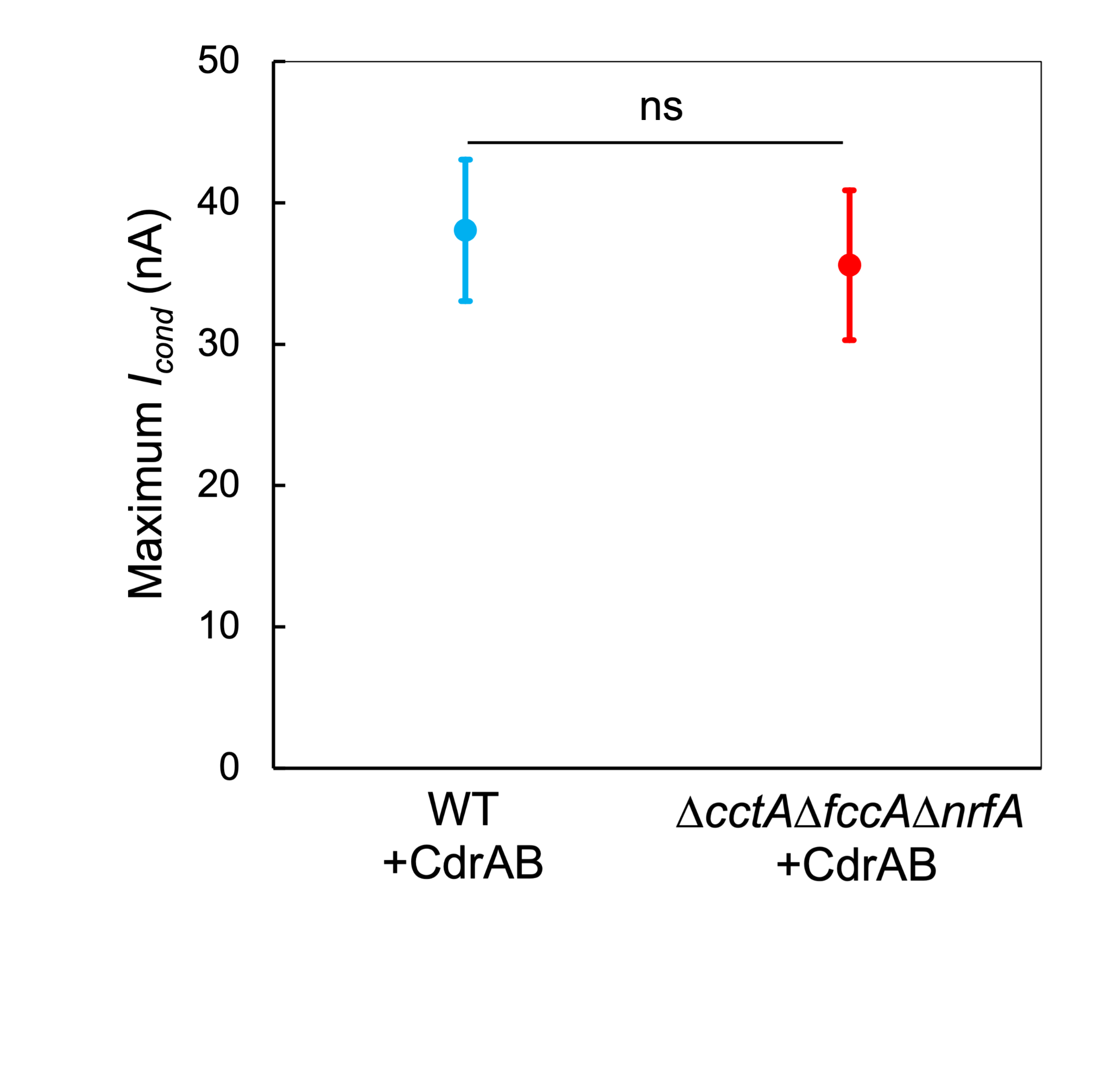

Figure S5. Maximum conduction currents of WT+CdrAB and ∆*cctA*∆*fccA*∆*nrfA*+CdrAB patterned biofilms from electrochemical gating measurements. The data (mean ± SD) were obtained from three biological replicates. The maximum conduction current is 38.04 ± 5.29 nA for the WT+CdrAB and 35.60 ± 5.30 nA for the ∆*cctA*∆*fccA*∆*nrfA*+CdrAB. No significant difference in maximum conduction current was observed between patterned WT+CdrAB biofilms and ∆*cctA*∆*fccA*∆*nrfA*+CdrAB biofilms (two-tailed unpaired *t*-test, *p* = 0.6021). Significance is indicated as ns (not significant, *p* > 0.05).

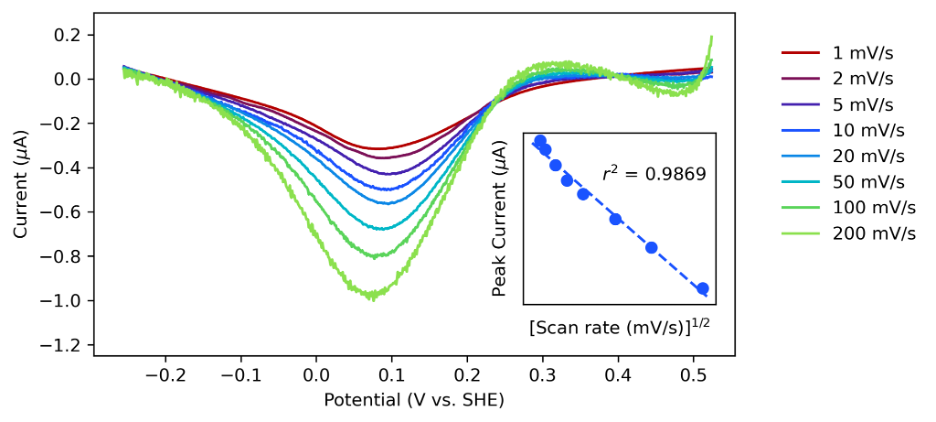

Figure S6. Scan rate dependent non-turnover cyclic voltammetry on WT+CdrAB biofilms patterned onto IDAs. Inset figure shows the peak currents observed in the data increase with increasing scan rate and clearly scale linearly with the square root of the scan rate.

Table S1. Strains and plasmids used in this study.

| **Strain** | **Description** | **Source** |
| --- | --- | --- |
| WT | *Shewanella oneidensis* MR-1, wild type strain | Lab stock |
| ∆*cctA*∆*fccA* (JG3107) | MR-1 derivative without genes encoding periplasmic cytochromes CctA and FccA | (2) |
| ∆*cctA*∆*fccA*∆*nrfA* (JG4347) | MR-1 derivative without genes encoding periplasmic cytochromes CctA, FccA, and NrfA | This work |
| WT+CdrAB | MR-1 with pDawn-CdrAB | (1) |
| ∆*cctA*∆*fccA*+CdrAB | JG3107 with pDawn-CdrAB | This work |
| ∆*cctA*∆*fccA*∆*nrfA*+CdrAB | JG4347 with pDawn-CdrAB | This work |
| **Plasmids** | **Description** | **Source** |
| pDawn-CdrAB | Expression of CdrAB in response to blue light using the pDawn genetic circuit | (1) |
